# A multi-omic atlas of the human frontal cortex for aging and Alzheimer's disease research

**DOI:** 10.1101/251967

**Authors:** Phillip L De Jager, Yiyi Ma, Cristin McCabe, Jishu Xu, Badri N. Vardarajan, Daniel Felsky, Hans-Ulrich Klein, Charles C. White, Mette A. Peters, Ben Lodgson, Parham Nejad, Anna Tang, Lara M. Mangravite, Lei Yu, Chris Gaiteri, Sara Mostafavi, Julie A. Schneider, David A. Bennett

## Abstract

We initiated the systematic profiling of the dorsolateral prefrontal cortex obtained from a subset of autopsied individuals enrolled in the Religious Orders Study (ROS) or the Rush Memory and Aging Project (MAP), which are jointly designed and belong to a very few prospective studies of aging and dementia with detailed, longitudinal cognitive phenotyping during life and a quantitative, structured neuropathologic examination after death for >3,322 subjects. Here, we outline the first generation of data including genome-wide genotypes (n=2,090), whole genome sequencing (n=1,179), DNA methylation (n=740), chromatin immunoprecipitation with sequencing using an anti-Histone 3 Lysine 9 acetylation (H3K9Ac) antibody (n=712), RNA sequencing (n=638), and miRNA profile (n=702). Generation of other omic data including ATACseq, proteomic and metabolomics profiles is ongoing. Thanks to its prospective design and recruitment of older, non-demented individuals, these data can be repurposed to investigate a large number of syndromic and quantitative neuroscience phenotypes. The many subjects that are cognitively non-impaired at death also offer insights into the biology of the human brain in older non-impaired individuals.

## Background & Summary

Alzheimer’s disease (AD) is an age-related neurodegenerative disease with continuously progressive deleterious reactions within neuronal cells. Prospective studies in humans are capable to provide unique and important information of the full landscape of the progression of disease, which play critical role for our understanding of the cause and treatment strategy. Our samples come from two prospective studies of aging – The Religious order Study (ROS) and the Memory and Aging Project (MAP) – that recruit older individuals without known dementia and include (1) detailed cognitive, neuroimaging and other ante-mortem phenotyping and (2) an autopsy at the time of death that includes a structured neuropathologic examination. Both studies are run by the same team of investigators at the Rush Alzheimer Disease Center (RADC), and they were designed to be used in joint analyses to maximize sample size. ROS subjects live in communities distributed throughout the U.S., while MAP subjects live in communities in the Chicago metropolitan area. We refer to the joint dataset as “ROSMAP”. Cross-sectional assessment of cognitive performance at the last clinical evaluation can be used in analyses with neuropathology or brain omics data; however, the trajectory of cognitive decline is a more pertinent trait for drug discovery as this is the clinical outcome of interest in the vast majority of clinical trials both in the preclinical and clinical AD space. The primary trait that captures this trajectory of decline is the “Global Cognitive Slope”. It is derived from the annual neuropsychologic evaluation of each subject. 19 different neuropsychologic tests are common between ROS and MAP (of the 21 tests deployed by one or the other study), and these data are collapsed into a single “Global Cognitive Score.” The longitudinal Global Cognitive Scores are then used in a random effects model to estimate person-specific annual rates of cognitive decline controlling for known confounders such as demographics and years of education. The approach used in constructing these traits is described in detail elsewhere^1,2^ and can be applied to specific cognitive domains, e.g., episodic memory decline, one of the hallmarks of AD. In addition, cognitive and pathologic data can be integrated to generate new traits that capture the difference between observed cognitive function and the extent of neuropathologies present in each individual’s brain: for example, previous studies have generated measures of residual cognitive decline^3^ or residual cognition, after accounting for a participant’s neuropathologic burden^4^.

Cataloguing the multi-omic data of the neuronal cells in all the ROSMAP subjects regardless of their disease trajectory can provide insight to the molecular events that contribute to aging-related cognitive decline. We generated complementary sets of data from the dorsolateral prefrontal cortex (DLPFC) of individuals in the study after their death. All available brains at the time of funding were used in each omic generation from the DLPFC. Selection of subjects for genome-wide genotyping was different as that was performed from all self-reported non-Latino whites. Whole genome sequencing was limited to subjects with autopsy data. The data described in this report represent data that exist today and are available on Synapse. Numerous additional layers of data, including proteomic and metaboloimc data, from tissue samples and purified cell populations are being produced and will become available as the data are finalized. We look forward to this large set of molecular and phenotypic data being repurposed by the neuroscience and other communities of researchers.

## Methods

### The Religious order Study and the Memory and Aging Project (ROSMAP)

The ROS and MAP cohorts have been designed for data and sample sharing, and they have been at the forefront of large-scale omic data generation from the human brain and also of sharing such data through efforts such as the DREAM challenge^5^ and the AMP-AD research program funded by the National Institute of Aging. Previous reports described the study design and data collection scheme of each study in detail^6,7^. By October 8, 2017, 3,322 ROSMAP participants (72.7% females) were enrolled and completed the baseline assessment, of which 1,702 (67.3% females) had died and 1,475 (67.2% females) had undergone brain autopsies. The autopsy rate in these studies exceeds 86%, ensuring that the autopsied subjects are representative of the study populations. **Tables 1** and **2** outline the demographic and selected diagnostic characteristics of the subjects included for each set of data; they also list the most commonly used phenotypes. **Figure 1** illustrates the extent of subject overlap among the different sets of data.

**Figure 1.**
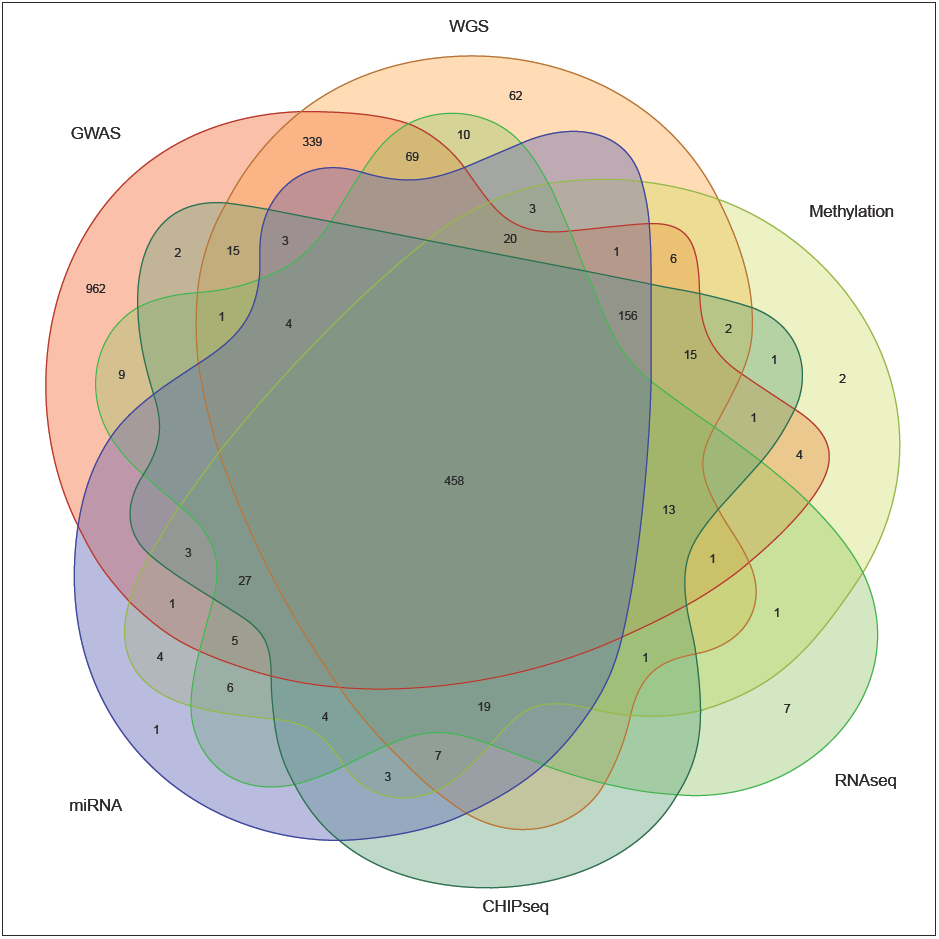
Overlap of the different layers of “omic” data. The venn diagram illustrates the extent to which the different layers of overlap in the ROS and MAP subjects that have been processed to date. 458 subjects have all layers of data described in this report.

**Table 1.** Demographic and diagnostic features of the ROS and MAP subjects in each layer of data^#^.

| Data type | N of all files | N of subjects with phenotypes | N of subjects with phenotypes on Synapse | mean age at death* | female (%) | AD (N) | MCI (N) | NCI (N) | other Dementia (N) |
| --- | --- | --- | --- | --- | --- | --- | --- | --- | --- |
| GWAS | 2090 | 2090 | 1036 | 86.7 (4.5) | 662 (63.9%) | 421 | 258 | 331 | 22 |
| WGS | 1196 | 1179 | 987 | 86.7 (4.4) | 638 (64.6%) | 405 | 245 | 313 | 21 |
| DNA methylation | 740 | 740 | 725 | 86.3 (4.7) | 460 (63.4%) | 305 | 172 | 229 | 19 |
| RNA-seq | 639 | 638 | 638 | 86.7 (4.5) | 408 (63.9%) | 254 | 169 | 201 | 12 |
| H3K9Ac-seq | 728 | 712 | 701 | 86.5 (4.6) | 454 (64.8%) | 293 | 171 | 219 | 18 |
| miRNA | 744 | 702 | 691 | 86.4 (4.6) | 443 (64.1%) | 290 | 165 | 219 | 17 |

<sup>#</sup>Data are based on the clinical data deposited on the Synapse and the age at death of >90+ were transferred to 90.
\*Values are presented in mean (standard deviation)
Abbreviation: AD, Alzheimer's disease; MCI, mild cognitive impairment; NCI cognitively non-impaired.

Phenotypic data is accruing continually in ROS and MAP, and new phenotypes are periodically added to the routine clinical and pathological data collection. Thus, these new phenotypes become available as additional neuropathologic and other characterizations are performed. The ante-mortem and neuropathologic traits that are currently available can be browsed to assemble biological sample sets and data sets with the features desired by the investigator through the RADC Research Resource Sharing Hub (https://www.radc.rush.edu/;jsessionid=8D1CD9FBE972335E0DD0494972E83B2A). The high-dimensional data described in this manuscript can be obtained through the Accelerating Medicnes Partnership for Alzheimer’s disease (AMP-AD) Knowledge Portal that is supported by the National Institute of Aging (https://www.synapse.org/ampad). The phenotypes listed in Table 2 are available through this portal, and additional phenotypic data are available from RADC. **Table 3** outlines the data through the portal. Additional data are being produced and will be available through the portal as they complete quality control analyses.

**Table 2.** List of traits available in Synapse for each subject.

| N | Traits | Description |
| --- | --- | --- |
| 1 | Age with the first diagnosis of AD | Float variable for age at cycle where first AD diagnosis was given. |
| 2 | Age at death | It is calculated from subtracting date of birth from date of death and dividing the difference by days per year (365.25). |
| 3 | Post-mortem interval in hours | Interval between death and tissue preservation in hours. |
| 4 | APOE genotype | Genotyping was performed by Agencourt Bioscience Corporation utilizing high-throughput sequencing of codon 112 (position 3937) and codon 158 (position 4075) of exon 4 of the APOE gene on chromosome 19. |
| 5 | Braak Stage | A semiquantitative measure of neurofibrillary tangles and the diagnosis includes algorithm and neuropathologist's opinion. |
| 6 | Diagnosis of AD by NIA-Reagan score | Diagnosis of AD by NIA-Reagan score. |
| 7 | The Mini Mental State Examination at the first diagnosis of AD. | A widely used, 30 item, standardized screening measure of dementia severity. |
| 8 | The Mini Mental State Examination at the last valid level. | A widely used, 30 item, standardized screening measure of dementia severity. |
| 9 | Assessment of neuritic plaques | A semiquantitative measure of neuritic plaques and the diagnosis includes algorithm and neuropathologist's opinion. |
| 10 | Final clinical consensus diagnosis | At the time of death, all available clinical data were reviewed by a neurologist with expertise in dementia, and a summary diagnostic opinion was rendered regarding the most likely clinical diagnosis at the time of death. Summary diagnoses were made blinded to all postmortem data. Case conferences including one or more neurologists and a neuropsychologist were used for consensus on selected cases. |

**Table 3.**
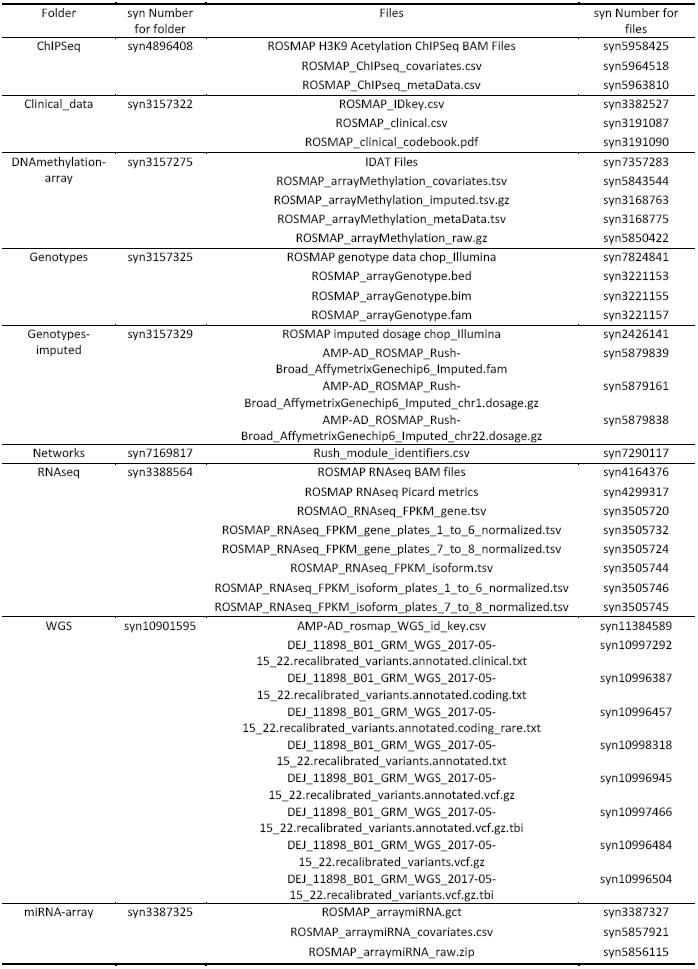
ROSMAP files deposited in AMPAD portal (syn3219045)

An important element of the design of the ROS and MAP studies is that all individuals are without known dementia at the time of entry into the study. Their cognitive trajectory is captured using a detailed battery of neuropsychological tests that is deployed annually to all living subjects. Subjects are also evaluated neurologically every year, and, at the time of death, a review of all ante-mortem data leads to a final clinical diagnosis for each participant: each individual receives a diagnosis of syndromic Alzheimer’s disease (AD), of mild cognitive impairment (MCI), or of being cognitively non-impaired (NCI). After the autopsy is concluded, a spectrum of neuropathologic diagnoses are obtained, such as a pathologic diagnosis of AD as defined using the modified NIA Reagan criteria^8^. However, many other pathologies are present in the brains of older individuals (the mean age of death is 88.8 years old in ROSMAP), and they are catalogued for each participant. As shown by **Figure 2**, there are imperfect overlaps of the two types of AD diagnoses. There are 95 participants with clinical AD without a pathological AD diagnosis, while 174 cognitively non-impaired participants have a pathologic diagnosis of AD.

**Figure 2.**
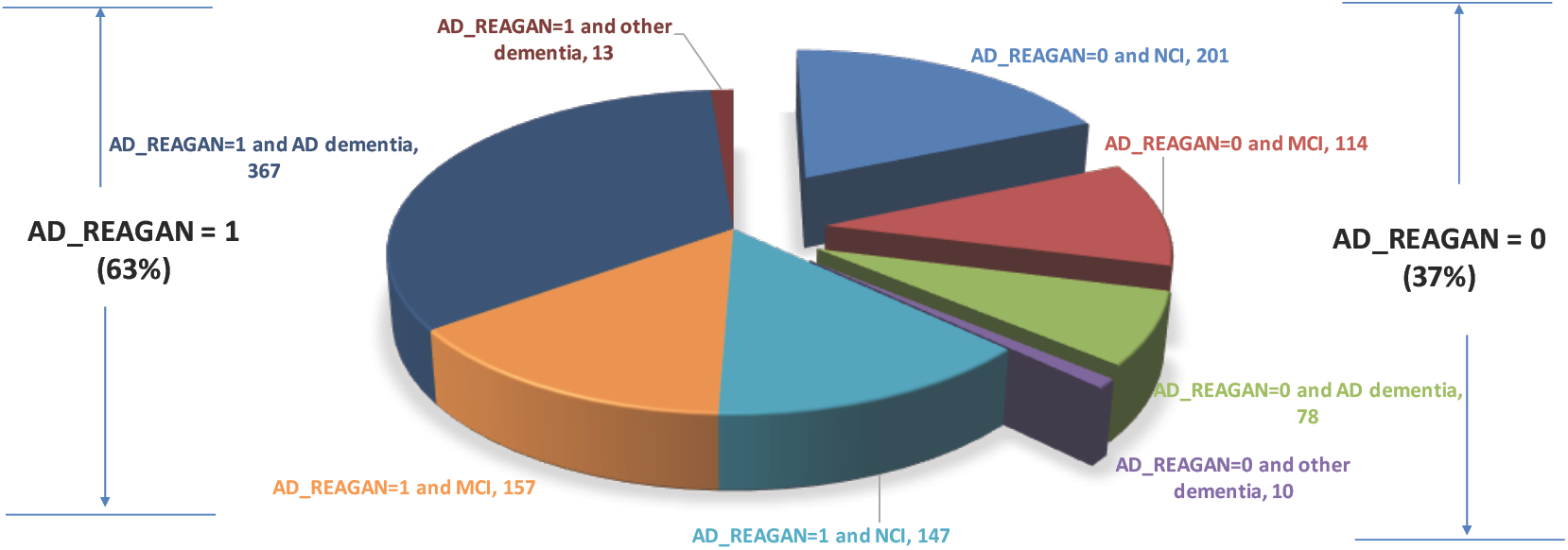
Overlaps between pathologic and clinical diagnosis of AD in ROSMAP.

### Molecular data generation

The RADC maintains a sample archive that contains DNA samples from each subject as well as the brains of deceased ROS subjects and the brain, spinal cord, selected muscles and nerves of MAP subjects. One hemisphere is cut into coronal slabs and frozen; the other hemisphere is fixed in 4% paraformaldehyde. Samples can be requested through the RADC website (www.radc.rush.edu).

#### Genotype data

DNA used for genotyping ROS and MAP participants was collected from postmortem brain tissue, whole blood, or lymphocytes. The majority of samples were genotyped on the Affymetrix GeneChip 6.0 platform (Santa Clara, CA, USA) at the Broad Institute’s Center for Genotyping (n=1204) or the Translational Genomics Research Institute (n=674). Additionally, 566 participants were genotyped on the Illumina OmniQuad Express platform at Children’s Hospital of Philadelphia. The same QC protocol was applied to all datasets using PLINK^9^ (pngu.mgh.harvard.edu/purcell/plink). We have limited analyses to participants of European decent. On the SNP level, we applied the following quality control (QC) filters: a genotype call rate > 95%, MAF > 0.01, misshap test 1x10^-9^, and a Hardy-Weinberg p < 0.001. The EIGENSTRAT software was used to calculate principle components used to control for population sub-structure; the top three principal components (PC)s are sufficient to correct for residual stratification.^10^

The final QC’ed dataset consists of 1709 participants of European ancestry from the Affy 6.0 platform and 384 participants from the Illumina platform. Using Beagle software (version: 3.3.2) and the 1000 Genomes Project (2011, Phase 1b data freeze) as a reference, dosage data was imputed on > 35 million SNPs for all genotyped samples who passed QC. We performed imputation separately for each genotyping platform. After removing SNPs with a MAF < 0.01 or an imputation quality info score < 0.3, approximately 7.5 million SNPs remained to analyze. Further information regarding genotyping and imputation can be found in previous publications.^11,12^

Following substantial improvements in phasing software and haplotype reference panels for populations of Caucasian ancestry, a second generation of imputation was performed for the autosomes in March, 2017 on the Michigan Imputation Server (MIS), using Minimac3, the Haplotype Reference Consortium (HRC) reference panel (v.1.1), and Eagle (v2.3) phasing software. Pre-imputation quality control identified 23 subjects from the Affy6.0 platform dataset (initial n=1709) and 3 subjects from the Illumina platform dataset (initial n=384) with high proportions of missing genotypes (>0.5) for at least one 20MB region of the genome, yielding final sample sizes of n=1686 and n=381 imputed using MIS. After imputation, these datasets were merged into a single n=2067 fileset. The number of variants imputed with high confidence (INFO score > 0.8) was >11.2 million, representing a large increase over the 1000 Genomes Phase 1 imputation dataset in the number of high quality variants available for analyses. Comparisons of subject-level genotype discordance for overlapping SNPs between the 1000 Genomes Phase 1 imputation, the MIS imputation, and whole genome sequencing (WGS) found an average discordance of 0.7% for MIS and 2.5% for 1000 genomes Phase 1 against WGS across all 22 chromosomes. This is a non-trivial increase in imputation quality and highlights nearly seven years of scientific improvement in the area of genomic imputation.

#### Whole Genome Sequence data

A subset of the ROSMAP samples (n=1200 for 1179 unique deceased participants) underwent whole genome sequencing, with DNA coming from brain tissue (n=806), whole blood (n=389) or lymphocytes transformed with EBV virus (n=5). WGS libraries were prepared using the KAPA Hyper Library Preparation Kit in accordance with the manufacturer’s instructions. Briefly, 1 ug of DNA was sheared using a Covaris LE220 sonicator (adaptive focused acoustics). DNA fragments underwent bead-based size selection and were subsequently end-repaired, adenylated, and ligated to Illumina sequencing adapters. Final libraries were evaluated using fluorescent-based assays including qPCR with the Universal KAPA Library Quanitification Kit and Fragment Analyzer (Advanced Analytics) or BioAnalyzer (Agilent 2100). Libraries were sequenced on an Illumina HiSeq X sequencer (v2.5 chemistry) using 2 x 150bp cycles.

Sequencing reads were aligned to the human reference using BWA-mem (version 0.7.15)^13^. Resulting BAM files contain all reads (passing or failing vendor quality checks), whether or not they aligned. Duplicate reads were detected and marked using Picard’s MarkDuplicates module (version 2.4.1) [http://picard.sourceforge.net]. Local alignment was performed around indels to identify putative insertions or deletions in the region using the GATK^14,15^ (version 3.5) indel realignment tool. Base quality score recalibration was performed using the GATK BQSR. This step uses observed data to improve the quality scores for each base in the sequence. GATK HaplotypeCaller and GenotypeGVCFs modules were used to generate individual genotype calls in genomic VCF and VCF format. Following variant calling, we ran the variant quality recalibration step in the GATK pipeline to empirically calibrate high quality variants. To ensure a high level of accuracy in genotype calls from sequencing, variants were filtered for minimum read depth (DP), variant calling confidence score (QD), VQSLOD and mapping and variant quality scores (MQ, GQ). Variant-level QC was performed using PLINK^9^ which includes checking genotype concordance using previous GWAS data, excluding variants with excess and/or systematic genotype missingness, examining departure from Hardy-Weinberg Equilibrium and identifying Mendelian inconsistencies among related individuals. Variants were annotated using ANNOVAR^16^. Variants were annotated with population frequencies in existing variant databases including dbSNP, 1000 Genomes, and the Exome Aggregation Consortium (ExAC). Prediction of variant function was obtained from POLYPHEN^17^ and SIFT^18^, cross-species conservation scores were obtained from PhyloP^19^,PhastCons^20^ and GERP^19^ and disease association were performed using OMIM^21^, HGMD^22^, ClinVar^23^.

#### RNA Sample Preparation

Approximately 100 mg of gray matter tissue from the dorsolateral prefrontal cortex (DLPFC) were sectioned while still frozen and shipped on dry ice overnight from the RADC to the Broad Institute. These sections were partially thawed on ice prior to dissection with a scalpel to separate the gray from the white matter and vasculature. Between 50mg and 100mg of gray matter was then added to 1ml of Trizol and homogenized with a 5mm stainless steel bead for 30 seconds at 30 Hertz using the Qiagen TissueLyser II. Following a quick spin to settle the foam, we would invert the tube 2-3 times to observe if the sample was fully homogenized. If chunks of tissue were still observed the sample was put back in the TissueLyser for another round. Homogenate was incubated at room temp for 5 minutes and then frozen at −80˚C. Samples were later thawed and processed in batches of 12-24 samples for RNA extraction using the Qiagen MiRNeasy Mini (cat no. 217004) protocol, including the optional DNAse digestion step. This protocol yields total RNA that includes miRNA. Samples were quantified by nanodrop and/or the RiboGreen assay; for each sample, the RNA Integrity Number (RIN) was measured using the Agilent Bioanalyzer Eukaryotic Total RNA Nano chip.

#### RNA Sequencing (RNA-Seq)

Samples were submitted to the Broad Institute’s Genomics Platform for transcriptome library construction following the dUTP protocol^24^ and Illumina sequencing. 5 micrograms of total RNA as measured by RiboGreen at a concentration of 50 nanogram/microliter with RNA Integrity Number (RIN) score of 5 or better were submitted for cDNA library construction. RIN score affects the fragment lengths of RNA inserts for library construction, and therefore we batched samples according to RIN scores so that library pools would have uniform insert sizes. 582 subjects in 6 batches/plates containing up to 92 samples, were processed using the dUTP method, barcoded and pooled for sequencing. Subsequently, 52 samples in a single batch were processed using the newer Illumina TruSeq method modified by The Broad Institute Genomics Platform to be strand specific and to use larger insert sizes. The resulting library closely resembles the library obtained by the dUTP method. The Truseq method uses only 250 nanograms of RNA input. Sequencing was carried out using the Illumina HiSeq2000 with 101bp paired end reads for a targeted coverage of 50M paired reads. The average sequencing depth was 50 million paired reads per sample. All reads were originally aligned by Tophat^25^ to the whole human genome reference (hg19) with Bowtie1 as aligner. Several Picard metrics (http://broadinstitute.github.io/picard) were collected from alignment results. Based on those Picard metrics, we implemented a paralleled and automatic RNA-Seq pipeline, in order to achieve higher quality of alignment and better estimation on gene expression levels. This pipeline includes identifying and trimming low quality bases (Q10) from beginning and end of each reads, identifying and trimming adapter sequencing from reads, detecting and removing rRNA reads and aligning reads to a transcriptome reference by a non-gap aligner (Bowtie1). The expression levels of gene and transcripts were estimated by RSEM package^26^. The Gencode V14 annotation were used by RSEM in the quantification process. Fragments Per Kilobase of transcript per Million mapped reads (FPKM) values were the final output of our RNASeq pipeline. 638 subjects passed QC from these two batches of samples.

Recently, the data were reprocessed in parallel with other AMP-AD RNAseq datasets, and this second version of the data is available as well. The input data for the RNAseq reprocessing effort was aligned reads in bam files that were converted to fastq using the Picard SamToFastq function. Fastq files were re-aligned to the GENCODE24 (GRCh38) reference genome using STAR with twopassMode set as Basic. Gene counts were computed for each sample by STAR by setting quantMode as GeneCounts, and transcript abundance estimated using Sailfish (see https://www.synapse.org/#!Synapse:syn9702085 for details)

#### miRNA profile

The RNA samples used to generate the RNAseq data were also submitted to the Broad Institute’s Genomics Platform for processing on the Nanostring nCounter platform to generate miRNA profiles for 800 miRNAs using the Human V2 miRNA codeset. The complete list of miRNAs is available at www.nanostring.com. 100 ng of each total RNA sample was used in the following Nanostring protocol: (1) multiplexed annealing of specific tags to their target miRNAs, (2) hybridization at 65°C for 16 h, (3) enzymatic purification to remove unligated material, (4) scanning for 600 fields of view on the nCounter Digital Analyzer. Raw data were normalized using the internal positive spike-in controls and the average counts of all endogenous miRNAs in each lane to account for the variability in both the hybridization process and sample input. A metric yielding a detection call at a confidence level of 95% (p<0.05) was determined.

The miRNA from the Nanostring RCC files were re-annotated to match the definitions from the miRBase v19. The raw data from the Nanostring RCC files were accumulated and the probe-specific backgrounds were adjusted according to the Nanostring guidelines with the corrections provided with the probe sets. After correcting for the probe-specific backgrounds, a three-step filtering of miRNA and sample expressions was performed. First, miRNA that had less than 95% of samples with a missing expression level were removed. This is followed by removing samples that had less than 95% of miRNAs with missing expressions. Thus, the call-rates for the samples and the miRNA are set at 95%. Finally, all miRNA whose absolute value is less than 15 in at least 50% of the samples were removed to eliminate miRNA that had negligible expression in brain samples. After the miRNA and sample filtering, the dataset consisted of 309 miRNAs and 702 subjects. A combination of quantile normalization and Combat^27^, specifying the cartridges as batches for the miRNA data, was used to normalize the data sets.

#### H3K9Ac ChIP-Seq

We identified the Millipore anti-H3K9Ac mAb (catalog # 06-942, lot: 31636) as a robust monoclonal antibody for our chromatin immunoprecipitation experiment. 50 milligrams of gray matter was dissected on ice from biopsies of the DLPFC of each ROS and MAP subject. The tissue was minced in a wash of ice cold PBS containing the Complete Protease Inhibitor Cocktail (Roche 11 836 170 001) and cross-linked with 1% formaldehyde at room temperature for 15mins and quenched with 0.125M Glycine. The tissue was then homogenized in cell lysis buffer (20mM Tris-HCl pH8.0, 85mM KCl, 0.5% NP 40) using the Tissue Lyser and a 5mm stainless steel bead. Then the nuclei were lysed in nucleus lysis buffer (10mM Tris-HCl, pH7.5, 1% NP 40, 0.5% sodium deoxycholate, 0.1% SDS) and chromatin was sheared using a Branson Sonifier 250 set to 40% amplitude for 0.7seconds on and 1.3 seconds off for 6 minutes with the thermal block set at −6˚C to generate the optimal majority fragment size range between 200 and 600bp. Samples were then centrifuged to pellet debris and 500ul of the supernatant – which is roughly half of the total volume – was incubated overnight at 4˚C with 2.5uL of the antibody with a final volume of 3mL using the ChIP Dilution Buffer (0.01% SDS, 1.1% Triton X-100, 1.2mM EDTA, 16.7mM Tris-HCl pH8.1, 167mM NaCl). Chromatin labeled with the H3K9Ac mark and bound to the antibody was purified with protein A sepharose beads, and the captured chromatin fragments were reverse cross-linked overnight in 250mM Tris-HCl, pH6.5, 62.5mM EDTA pH8.0, 1,25M NaCl, 5mg/mL Proteinase K, 62.5 ug RNaseA at 65˚C. The captured DNA fragments were then extracted using a phenol:chloroform phase separation, and prepared for library construction using the END-IT DNA repair kit (Epicenter Cat. No. ER0720), and single 3’-adenine overhangs were added using Klenow(3’-5’ exo-) (New England Biolabs, Cat. No. M0212L). Qiagen MiniElute spin columns were used to clean up each of these reactions. Barcoded Illumina adapters prepared by the Broad Institute’s Genomic’s platform, were ligated to cleaned DNA fragments with DNA ligase (New England Biolabs, Cat. No. M2200S) and subsequently cleaned using 0.7X AMPure XP beads with 70% freshly prepared ethanol washes two times. The libraries were then amplified by PCR using PFU ULTRA II HS 2X Master Mix (Agilent Cat. No. 600852). Size selection was carried out by cutting the section between 275bp-600bp after running electrophoresis on a 2% agarose gel using a 100bp ladder (NEB-N3231S). The final library was extracted from the excised gel fragments using the Gel Extraction Kit (Qiagen-28706). Libraries were quantified by Qubit in triplicate and pooled for sequencing in 4-plex or 8-plex and sequenced for 36bp single end reads on Illumina’s HiSeq2000 splitting the two cohorts across the pools as evenly as possible and targeting about 20M reads per sample.

To quantify histone acetylation, after sequencing, single-end reads were aligned to the GRCh37 reference genome by the BWA algorithm, and duplicated reads were flagged using picard tools. Reads mapping to multiple locations were marked by setting the mapping quality to 0 and were excluded from subsequent processing. Peaks were detected by MACS2 using the option for broad peaks and a stringent q-value cutoff of 0.001. Pooled DNA of 7 samples was used as negative control. A combination of five ChIP-seq quality measures were employed to detect low quality samples: samples that did not reach (i) ≥ 15 10^6^ uniquely mapped unique reads, (ii) non-redundant fraction ≥ 0.3, (iii) cross correlation ≥ 0.03, (iv) fraction of reads in peaks ≥ 0.05 and (v) ≥ 6000 peaks were removed. Samples passing quality control were used to define a common set of peaks termed H3K9Ac domains. Each base overlapped by a peak in at least 100 samples (∼15%) was considered as part of an H3K9Ac domain. Domains within 100 bp distance were merged. Subsequently, H3K9Ac domains of less than 100 bp width were removed resulting in a total of 26,384 H3K9Ac domains with a median width of 2,829 bp. Finally, uniquely mapped unique reads were extended towards their 3’-end to the estimated fragment size, and the number of reads overlapping each domain was computed for each sample. In total, read count data and bam files are available for 712 subjects.

#### DNA Methylation profile

As was done in the RNA extraction effort, gray matter was dissected from white matter while on ice from a sample of frozen DLPFC. This cortical sample was then processed using the Qiagen QIAamp mini protocol (Part number 51306) to extract DNA. Samples were evaporated to increase concentration to 50ng/ul and submitted to the Broad Institute’s Genomics Platform for processing on the Illumina Infinium HumanMethylation450 BeadChip^28^.

Because of the use of different thermocyclers during data generation process, a strong batch effect was observed and we applied a series of strategies of quality control and data analysis to counter such batch effect. On the probe level QC, at first, we selected good quality probes according to the detection *P* value < 0.01 across all samples. We further removed those probes predicted to cross-hybridize with the sex chromosomes ^29^ and those having overlaps with known SNP with MAF 0.01 (±10bp) based on the 1000 Genomes database. On the subject level QC, we at first used principal component analysis (PCA) based on 50,000 randomly selected probes to select subjects that were within ±3 s.d. from the mean of a principal component (PC) for PC1, PC2, and PC3. Secondly, we filtered out those subjects with poor bisulfite conversion efficiency. We have compared data normalization strategy of COMBAT ^27^ and independent component analysis (ICA) (http://cran.r-project.org/web/packages/fastICA/index.html) with the adjustment of batch variable in the analysis, and we found that the adjustment of the batch variable outperform the other two strategies.

β values reported by the Illumina platform were used as the measurement of methylation level for each CpG probe tagged on the chip. We imputed those missing β values using a *k*-nearest neighbor algorithm for *k* = 100. The primary data analysis includes adjustment of age, sex, and experiment batch variable. We estimated the proportion of NeuN+ cells (primarily neurons) in each brain sample using DNA methylation data ^30^, but we did not find it had significant associations with a pathologic diagnosis of AD (*P* = 0.08). Overall, we have methylation profiles for 740 subjects.

## Data Records

For high-dimensional data, the NIA-supported AMP-AD Knowledge Portal on the Synapse platform is the preferred distributor (Data Citation 1), and additional samples as well as phenotypic and other data are available through the RADC Research Resource Sharing Hub (Data Citation 2). Data from each unique participant is assigned the same 6 digit study ID, facilitating the relation of different data types. To download files see the ‘How to Download’ guide on the folder to download all folder content, and the Synapse documentation for more details: http://docs.synapse.org/articles/downloading_data.html. The following are the key files: (1) Study description (Data Citation 3), (2) Clinical data, codebook and assay ID key (Data Citation 4), (3) ChIPseq (Data Citation 5), (4) DNA methylation (Data Citation 6), (5) Genotypes (Data Citation 7), (6) Genotypes imputed (Data Citation 8), (7) RNAseq (Data Citation 9), (8) miRNA: (Data Citation 10).

## Technical Validation

### Technical Validation

#### Data derived based on DNA: Genotype, imputation, whole genome sequence, and methylation

All DNA samples go through the same rigorous quality control process before and after genotype generation, so we see no difference in data quality based on source of DNA. Affymetrix GeneChip 6.0 platform and Illumina OmniQuad Express platform are well validated platforms for genotyping. Detailed QC pipeline was described in ^12^. Briefly, the standard QC measures for SNPs (HWE p > 0.001; MAF > 0.01; genotype call rate > 0.95; misshap test × 1 10-9) and subjects (genotype success rate > 0.95; genotype-derived gender concordant with reported gender, excess inter/intra-heterozygosity) were applied. The top hits of the genotype data were successfully replicated in another independent dataset ^12^. For the whole genome sequencing data, base quality score recalibration was performed using the GATK BQSR and the empirical calibration of the variant quality was done using GATK pipeline. Variants were further filtered for minimum read depth (DP), variant calling confidence score (QD), VQSLOD and mapping and variant quality scores (MQ, GQ). For the methylation data, we applied both probe-level (detection P ≥ 0.01 across all samples; not cross-hybridize with the sex chromosomes; not overlapped with known SNPs with MAF ≥ 0.01 within 10bp region) and subject-level QC (within 3 s.d. from the mean of a principal component (PC) for PC1, PC2 and PC3, and those with poor bisulfite conversion efficiency). The top hits were also successfully replicated in an independent sample ^28^. Experimental replicates and controls were designed to calibrate data.

#### Data derived based RNA: RNAseq and miRNA

The RNA extraction protocol, Qiagen MiRNeasy Mini (cat no. 217004) protocol, has been validated to be effective to purify both total RNA and miRNA ^31–36^. For each sample, the RIN score was measured using the Agilent Bioanalyzer Eukaryotic Total RNA Nano chip. Those RNA samples with RIN score of 5 or better were submitted for cDNA library construction. RIN score affects the fragment lengths of RNA inserts for library construction, and therefore we batched samples according to RIN scores so that library pools would have uniform insert sizes. In order to get correct alignment, we trimmed the reads if they have low quality bases (Q10) from beginning and end or those reads of adapters and rRNA. Experimental replicates and controls were designed to calibrate data.

#### H3K9Ac ChIP-Seq

Pooled DNA of 7 samples was used as negative control. A combination of five ChIP-seq quality measures were employed to detect low quality samples: samples that did not reach (i) ≥ 15 × 10^6^ uniquely mapped unique reads, (ii) non-redundant fraction ≥ 0.3, (iii) cross correlation ≥ 0.03, (iv) fraction of reads in peaks ≥ 0.05 and (v) ≥ 6000 peaks were removed.

## Usage Notes

All data are publically available following the completion of a data use agreement that can be completed through the RUSH University ADC (http://www.radc.rush.edu/requests.htm;jsessionid=A2587E2977951F15E7AEB2FF3812F66A) or the Synapse platform (https://www.synapse.org/#!AccessRequirements:ID=syn3219045&TYPE=ENTITY).

The ROS and MAP cohorts have useful features that allow the data generated from their subjects to be repurposed for many different analyses and to render results relevant to the population of older individuals. Most importantly, all subjects are community-dwelling without known dementia at the time of enrollment. All testing is performed in the participants’ homes, and the only inclusion criteria are age and willingness to sign the informed consent and Anatomical Gift Act. Thus, participants capture the full spectrum of phenotypes found in an aging human population. Further, both ROS and MAP include longitudinal rigorous clinical, functional, neuropsychologic and magnetic resonance imaging characterization of participants while they are alive, as well as a structured clinical and quantitative neuropathologic assessment at autopsy. The application of standard clinical scales to the collected data provide both syndromic diagnoses and semi-quantitative measures such as the many cognitive function tests that allow the comparison of results from ROS and MAP to those from other collections of subjects. These simpler phenotypes also enables us to contribute data to consortia for joint or meta-analyses, as has been done for a wide range of clinical, imaging and pathologic phenotypes^37–39^. As clinical and pathologic phenotypes do not occur in isolation, the deep clinical and neuropathologic phenotyping of each subject enables investigators to resolve the contribution of a given molecular feature to multiple different intermediate traits that ultimately contribute to cognitive decline and other common conditions of aging. We also note that certain limitations must be taken into account when interpreting results from these cohorts. (1) These cohorts sample a large spectrum of the older population but are not a random sample of the overall population; nonetheless, they capture a much larger spectrum of the aging population than most autopsy series that rely on the subset of individuals coming to the attention of the health care system because of their symptoms and often have highly selective recruitment criteria. (2) The mean age at study entry is 78.9 (SD=7.5, range 55.4-102.1), and the mean age at death is 89 (SD=6.6, range 65.9-108.3). Since subjects are older and without known demented at study entry, there is a bias in study entry stemming both from survival to older age from all causes of early mortality and from surviving to study entry without significant cognitive impairment. (3) The range of age at the time of death is broad but restricted to the older segment of the age distribution of the North American population. (4) Finally, agreement for organ donation likely introduces a subtle bias. However, it should be noted essentially all risk factors for AD dementia identified in the cohort have been replicated in other cohort studies.

We also note that many of these neuropsychologic and neuropathologic traits are correlated (**Table 4**) and that many of these traits correlate with advancing age. The age and sex of subjects have very strong effects on the brain’s epigenome and transcriptome; these two variables are important confounders when performing any analyses of ROS and MAP data. Further, one must carefully consider the molecular effects of neuropathologies that confound aging-related analyses as we have shown with the methylome^40^. Finally, both circadian and seasonal rhythms influence the epigenome and the transcriptome, introducing an important source of variation for many genes that is rarely appreciated^41^.

**Table 4.** Correlation matrix of cognitive traits with age at death.

|  | Braak score | CERAD score | Mini-mental state exam | Age at death |
| --- | --- | --- | --- | --- |
| Braak score | 1.0 | -0.4<br>($P=6.7\times 10^{-47}$) | -0.6<br>($P=9.3\times 10^{-98}$) | 0.3<br>($P=4.3\times 10^{-27}$) |
| CERAD score | | 1.0 | 0.4<br>($P=6.7\times 10^{-35}$) | -0.2<br>($P=5.1\times 10^{-10}$) |
| Mini-mental state exam | | | 1.0 | -0.2<br>($P=9.6\times 10^{-10}$) |
| Age at death |  |  |  | 1.0 |

The ROS and MAP cohorts have been designed for data and sample sharing, and they have been at the forefront of large-scale omic data generation from the human brain and also of sharing such data through efforts such as the DREAM challenge^5^ and the AMP-AD research program funded by the National Institute of Aging. The data described in this report represent data that exist today and are available on Synapse. Numerous additional layers of data, including proteomic and metabolomic data, from tissue samples and purified cell populations are being produced and will become available as the data are finalized. We look forward to this large set of molecular and phenotypic data being repurposed by the neuroscience and other communities of researchers.

## Data Citations

1. https://www.synapse.org/#!Synapse:syn2580853 (2016).
2. AMP-AD Knowledge Portal. *Synpase Bionetworks* RADC Research Resource Sharing Hub https://www.radc.rush.edu/;jsessionid=8D1CD9FBE972335E0DD0494972E83B2A (2016)
3. *Sage Bionetworks* https://dx.doi.org/10.7303/syn321 (2015)
4. *Sage Bionetworks* https://dx.doi.org/10.7303/syn3157322 (2015)
5. *Sage Bionetworks* https://dx.doi.org/10.7303/syn4896408 (2015)
6. *Sage Bionetworks* https://dx.doi.org/10.7303/syn3157275 (2015)
7. *Sage Bionetworks* https://dx.doi.org/10.7303/syn3157325 (2015)
8. *Sage Bionetworks* https://dx.doi.org/10.7303/syn3157329 (2015)
9. *Sage Bionetworks* https://dx.doi.org/10.7303/syn3388564 (2015)
10. *Sage Bionetworks* https://dx.doi.org/10.7303/syn3387325 (2015)

## Acknowledgements

We are grateful to the participants in the Religious Order Study, the Memory and Aging Project. This work is supported by the US National Institutes of Health [U01 AG046152, R01 AG043617, R01 AG042210, R01 AG036042, R01 AG036836, R01 AG032990, R01 AG18023, RC2 AG036547, P50 AG016574, U01 ES017155, KL2 RR024151, K25 AG041906-01, R01 AG30146, P30 AG10161, R01 AG17917, R01 AG15819, K08 AG034290, P30 AG10161 and R01 AG11101].

